# Extracellular mechanical forces drive endocardial cell volume decrease during cardiac valve morphogenesis

**DOI:** 10.1101/2021.07.27.453460

**Authors:** Hélène Vignes, Christina Vagena-Pantoula, Mangal Prakash, Caren Norden, Florian Jug, Julien Vermot

## Abstract

Organ morphogenesis involves dynamic changes of tissue properties at the cellular scale. In addition, cells need to adapt to their mechanical environment through mechanosensitive pathways. How mechanical cues influence cell behaviors during morphogenesis, however, remains poorly understood. Here we studied the influence of mechanical forces during the formation of the atrioventricular canal (AVC) where cardiac valves develop. We show that in zebrafish the AVC forms within a zone of tissue convergence between the atrium and the ventricle which is associated with increased activation of the actomyosin meshwork and endocardial cell orientation changes. We demonstrate that tissue convergence occurs with a major reduction of endocardial cell volume triggered by mechanical forces and the mechanosensitive channels TRPP2/TRPV4. In addition, we show that the extracellular matrix component hyaluronic acid controls cell volume changes. Together, our data suggest that cell volume change is a key cellular feature activated by mechanical forces during cardiovascular morphogenesis. This work further unravels how mechanical forces and extracellular matrix can influence tissue remodeling in developing organs.

## INTRODUCTION

Most organs need to acquire a defined shape for optimal function. Organogenesis is controlled both spatially and temporally as a result of the interplay between genetic and mechanical factors (Mammoto & Ingber, 2010; Sivakumar & Kurpios, 2018). Mechanical cues can originate from diverse contexts: the cell environment for example the surrounding extracellular matrix (ECM), neighboring cells, or neighboring tissues (Charras & Sahai, 2014; Hannezo & Heisenberg, 2019; Martino et al., 2018; Petridou et al., 2017; Villedieu et al., 2020). Further, mechanical forces can originate from the tissue itself (Boselli et al., 2015; Xiong et al., 2020). Cells constitute the functional unit of every tissue and can sense and react to those different forces driving coordinated cell behaviors (cell shape changes, cell intercalation, cell apoptosis, etc) that are essential for tissue remodeling (Heisenberg & Bellaïche, 2013; Lecuit et al., 2011). Yet how mechanical forces affect cell behaviors and tissue shape at the cellular scale remains unknown.

Amongst the different cellular mechanisms that control morphogenetic processes (Heisenberg & Bellaïche, 2013; Lecuit et al., 2011; Mao & Baum, 2015), cell volume changes have recently emerged as an essential modulator of tissue shape. In *Drosophila Melanogaster*, cell volume decrease drives the tissue movement during dorsal closure, (Saias et al., 2015). Moreover, cell volume changes are also key in regulating the lumen volume of intestinal organoids (Yang et al., 2020) and the Küpffer vesicle morphogenesis in zebrafish (Dasgupta et al., 2018). *In vitro*, external mechanical cues can trigger cell volume changes (Guo et al., 2017; Wang et al., 2020; Xie et al., 2018). Cell volume is controlled by osmotic regulation, cell cytoskeletal contractility as well as cell growth and division (Cadart et al., 2019). However, how cell volume changes can lead to tissue remodeling in broader tissular contexts, such as the cardiovascular system, remains unclear.

The heart acquires its function early during embryonic development in order to pump blood throughout the body. Defects in heart formation and especially heart valve anomalies can lead to congenital heart diseases with major medical implications (Lincoln & Yutzey, 2011). The fluid shear stress and stretching forces generated by heartbeat have been reported to be essential for proper valve morphogenesis (Auman et al., 2007; Bartman et al., 2004; Hove et al., 2003; Kalogirou et al., 2014; Vermot et al., 2009). The inner tissue layer of the heart corresponds to the endocardium, constituted of specialized endothelial cells called endocardial cells (EdCs). Like endothelial cells, EdCs are potent mechanosensors and mechanotransducers of mechanical forces (Campinho et al., 2020). The membrane-bound stretch-sensitive channels Piezo1, TRPP2, and TRPV4 are important for mechanical force sensing during cardiac valve development (Duchemin et al., 2019; Faucherre et al., 2020; Heckel et al., 2015). Moreover, EdCs interact also with their ECM (i.e. cardiac jelly), produced by both myocardial and endocardial cells, and which plays key roles during cardiac morphogenesis (Steed et al., 2016; Derrick et al., 2019; Derrick & Noël, 2021; Grassini et al., 2018).

One of the main components of the cardiac jelly is hyaluronic acid (HA), which is a negatively charged glycosaminoglycan (GAG). HA has the property to generate osmotic pressure and therefore attracts water, resulting in a local swelling of the cardiac jelly at the AVC (Camenisch et al., 2000; Cowman et al., 2015; Lockhart et al., 2011; Schroeder et al., 2003; Tong et al., 2014). However, the roles of the forces generated by the heartbeat as well as the impact of biophysical properties of the ECM on the modulation of EdC shape during primitive heart tube formation are currently not well understood.

In zebrafish, the formation of a local cell cluster within the single-layered endocardial sheet at the AVC marks the onset of cardiac valve formation (Pestel et al., 2016; Steed et al., 2016). At the tissue level, this event is preceded by a symmetry-breaking event promoted by a tissue convergence towards the AVC (Boselli et al., 2017). Here, we used this morphogenetic model to investigate the early cellular features involved in endocardial tissue convergence, starting from 28hpf, when the zebrafish heart has a tubular structure, until 48hpf when the heart has looped and the two chambers (atrium, ventricle) are formed. We found that tissue convergence is associated with a global EdC orientation change directed towards the AVC and a cell clustering event that is associated with a local cell volume decrease of the AVC cells. Interestingly, neither the cell cluster formation nor the cell volume changes were linked to cell proliferation. At the molecular level, we found that the cell volume decrease depends on the mechanosensitive TRP channels (both TRPP2 and TRPV4) and the ECM component hyaluronic acid. We propose a model where mechanotransduction and the osmotic pressure generated by HA accumulating within the cardiac jelly dictates local cell volume changes in the endocardium. This may be a general feature by which mechanical forces shape the cardiovascular system, such as the heart or blood and lymphatic vessels in the vascular system.

## RESULTS

### EdC polarisation during AVC development

Using zebrafish for high precision live imaging, we analyzed the morphogenetic patterns of the endocardium at cellular resolution. Axial cell polarity is a well-established readout of the orientation and the coordination of cell movements within endothelial tissues (Franco et al., 2015; Kwon et al., 2016; Pouthas et al., 2008). Taking advantage of this fact, we investigated the dynamics of endocardial cell orientation during AVC development *in vivo* to characterize the general morphogenetic features underlying AVC formation. We used a zebrafish transgenic line that labels the Golgi apparatus and the nucleus specifically in endothelial cells *Tg(fli1:nEGFP); Tg(fli1a:B4GALT1-mCherry)* (Kwon et al., 2016), thereby providing a readout of the global tissue patterning at the cellular scale. Nucleus-to-Golgi axis orientation was analyzed every two hours from 28hpf to 36hpf and at 48hpf in order to track EdC orientation changes (Figure 1.A, Figure S1.A, Video S1). We classified the cells into three categories: 1) nucleus-to-Golgi axis towards the outflow, 2) nucleus-to-Golgi axis towards the inflow and 3) no clear nucleus-to-Golgi axis orientation (Figure 1.B; Figure S1.B). For statistical analysis, we quantified nucleus-to-Golgi axis changes in the ventricle (Figure 1.C) and in the atrium over time (Figure 1.C’). Before heartbeat (22hpf), 48.6±5.7% of the cells showed a nucleus-to-Golgi axis towards the outflow (n=130 cells, N=3 embryos) (Figure S1.C). After blood flow initiation, we found that the majority of the cells (66.5±2.0%, n=271 cells, N=5 embryos for the atrium and 70.2±4.6%, n=198 cells, N=5 embryos for the ventricle) showed a nucleus-to-Golgi axis towards the outflow at 28 hpf in both chambers (Figure 1 C-C’). While nucleus-to-Golgi axis patterns remained unchanged from 28hpf to 48hpf within the atrium (Figure 1.C’), ventricular cells gradually reversed their nucleus-to-Golgi between 28 and 48hpf (48.2±2.0%, n=578 cells, N=7 embryos) to point towards the AVC (Figure 1.C). These results indicate that tissue convergence is accompanied by a global orientation of the nucleus-to-Golgi axis towards the AVC, starting from 30hpf underlining the collective movements of the endocardial cells required to initiate the formation of cardiac valves (Figure 1.F).

**Figure 1.**
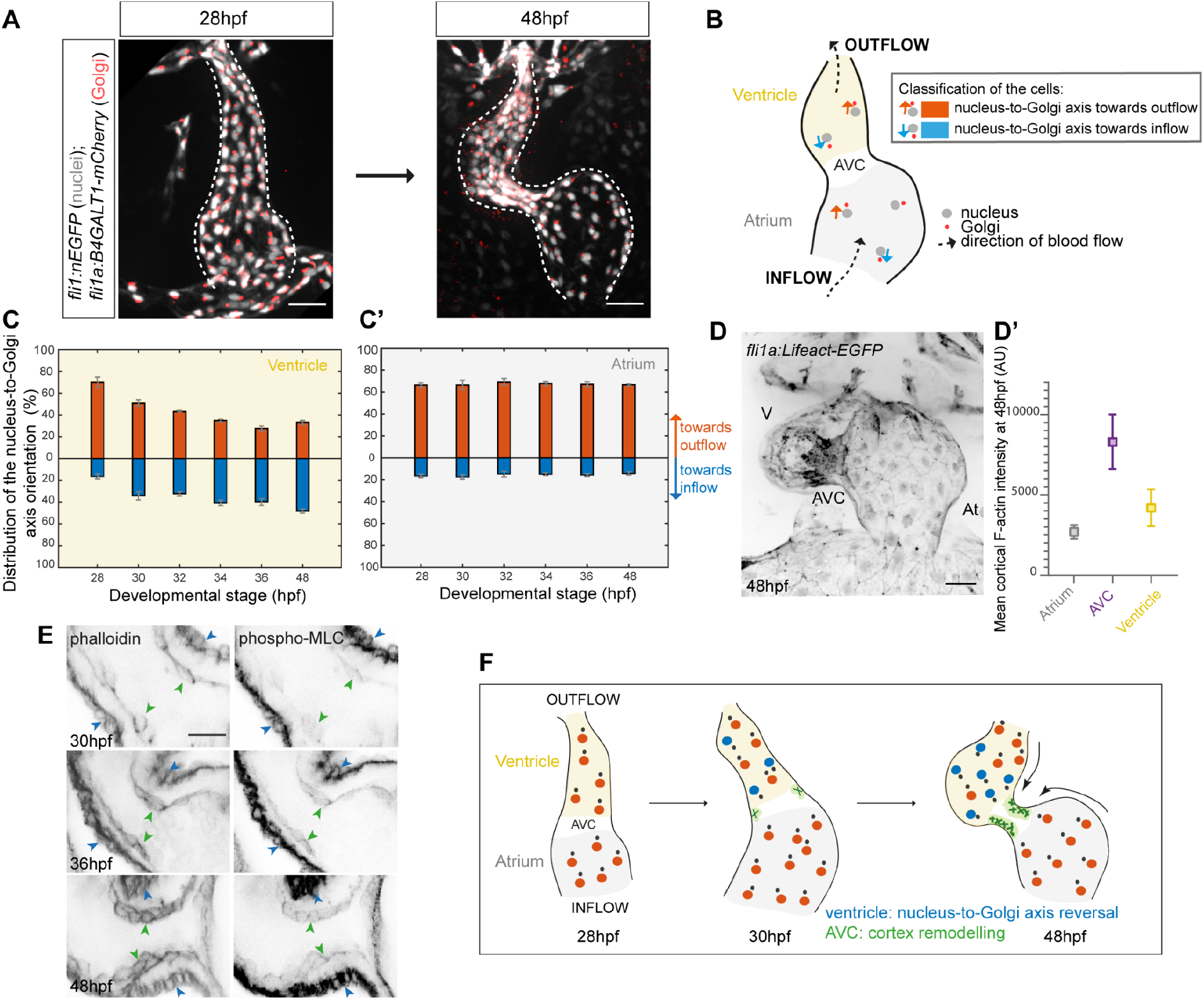
Ventricular EdC nucleus-to-Golgi axis reversal and F-actin remodeling at the AVC happens concomitantly from 30hpf to 48hpf. (**A**) Maximal projection of 28hpf to 48hpf hearts from *fli1a: B4GALT1-mCherry; fli1: nEGFP* embryos to track changes in EdC nucleus-to-Golgi axis orientation. White dotted lines outline the endocardium. Scale bars, 30μm. (**B**) Schematic explanation of the way cell orientation was classified within the endocardium. (**C-C’**) Quantification of the percentage of cells with the nucleus-to-Golgi axis oriented towards the outflow and towards the inflow from 28hpf to 48hpf (N=5 embryos, 28hpf ; N=5 embryos, 30hpf ; N=4 embryos, 32hpf ; N=4 embryos, 34hpf ; N=5 embryos, 36hpf ; N=7 embryos, 48 hpf ; Error bars show the s.e.m) in the ventricle (**C**) and in the atrium (**C’**). (**D**) Maximal projection of *fli1a: LifeAct-EGFP* heart showing an increase in F-actin fluorescence intensity at the AVC. V=ventricle, At= atrium. Scale bar, 30μm. (**D’**) Quantification of the F-actin fluorescent intensity based on the data in (D). A.U= arbitrary units. Error bars represent the s.d. (**E**) Single z-plane immunofluorescence with phalloidin and phospho-myosin light chain (MLC) antibody in the AVC showing the presence of a strong F-actin and phospho-MLC signal in the myocardium (blue arrows) and an increasing signal through time in the AVC (green arrows) from 30hpf, 36hpf, to 48hpf. Scale bars, 20μm (**F**) Schematic representation of the general cellular morphogenetic features underlying AVC development.

### EdC volume during AVC development

Considering the global change in EdC orientation at the onset of AVC morphogenesis, we reasoned that the cells located in the area of convergence had to specifically change shape within the AVC. We therefore hypothesized that a local cell size change at the AVC could contribute to the process of AVC morphogenesis. To address this, we analyzed EdC volume changes during AVC morphogenesis. We first quantified the cell area and the cell height (apicobasal direction) separately in order to compare the cell size between all three regions (atrium, AVC, ventricle) in order to calculate the mean cell volume. To measure the area of the apical cell surface, we used a transgenic zebrafish line (*TgBAC(ve-cad:ve-cad-TS)*) labeling vascular endothelial (VE)-cadherin (Lagendijk et al., 2017) (Figure 2.A). We estimated the cell size by multiplying the apical cell surface area by the cell thickness (Figure 2.B). Based on these calculations, the size of AVC cells is 412.2±21.6 µm^3^ (n= 41 cells), while that of cells located in the atrium and the ventricle are 1348±67.9 µm^3^ (n= 48 cells) and 933.1±47.8 µm^3^ (n= 34 cells), respectively (Figure 2.C). We concluded that cell size varies depending on the different regions of the heart with cells being smaller in the AVC.

**Figure 2.**
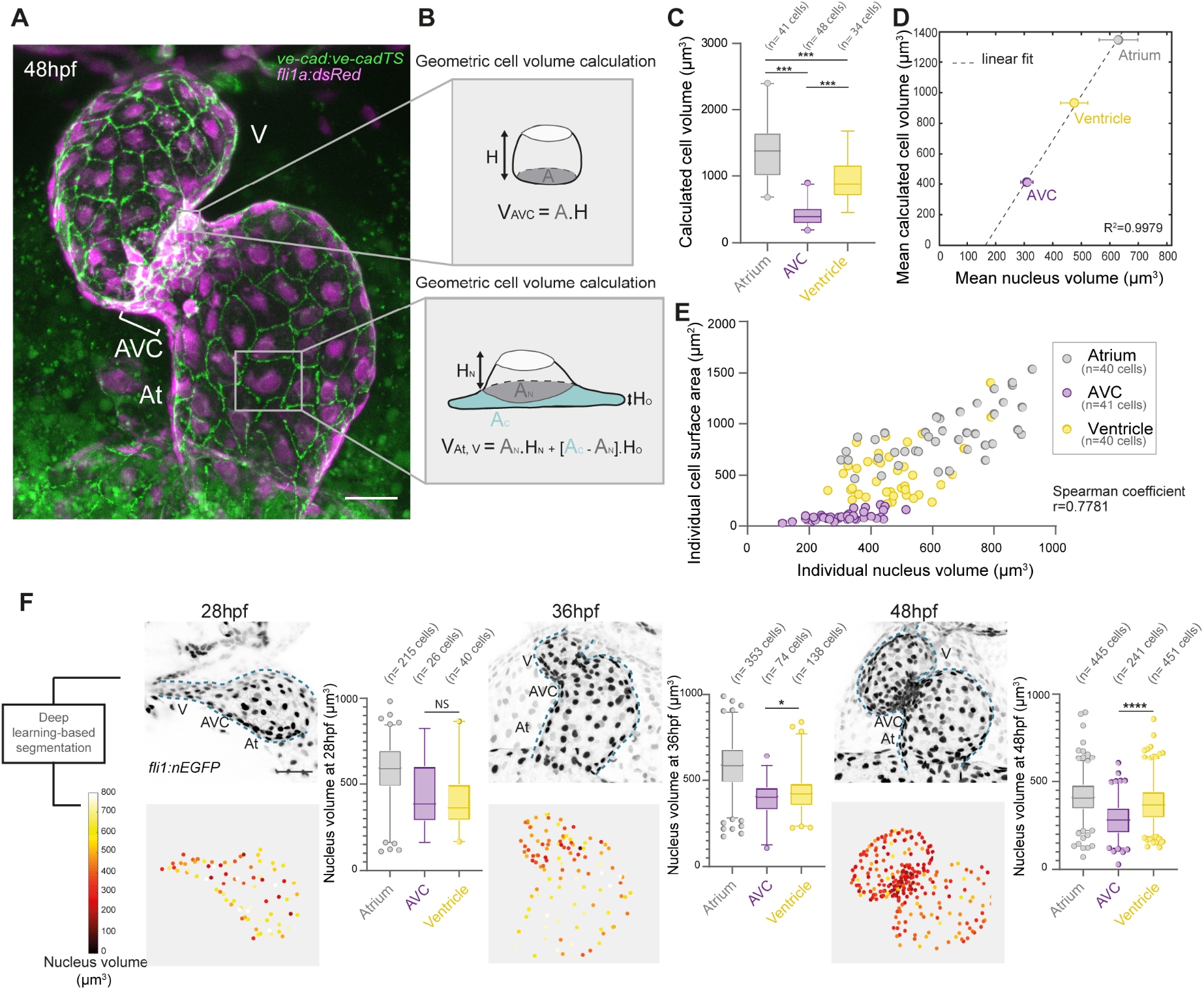
3D analysis of EdC size reveals a significant decrease in cell volume at the AVC. (**A**) Maximal projection of a stopped heart from a *ve-cad:ve-cadTS* ; *fli1a :dsRed* embryo. V=ventricle, At=atrium. Scale bar, 30 μm. (**B**) Schematics depicting methods used to calculate the cell volume in EdCs of the AVC (top panel) and in EdCs of the ventricle and the atrium (bottom panel). A= area, AN= area of the nucleus, AC= area of the cytoplasm, H= height of the cell, HN= Height at cell region containing the nucleus, HO= Height in other regions of the cell (**C**) Quantification of the cell volume shows differences between EdCs located in the atrium (n=41 cells), AVC (n=48 cells), and ventricle (n=34 cells) with AVC EdCs having the lowest cell volume. Mann-Whitney test. (**D**) Nucleus volume scales with cell volume. Correlation (R^2^=0.9979) between the mean cell volume (calculated in (C)) and the mean nucleus volume quantified via segmenting nuclei using deep learning. Error bars represent the s.e.m for both axes (**E**) Individual cell surface area is positively correlated with individual nucleus volume (Spearman coefficient= 0.7781). Each dot represents a cell analyzed for both its surface area and nucleus volume (**F**) Quantification of the nucleus volume at 28, 36 and 48hpf reveals a significant decrease in cell volume of the AVC EdCs over development time. Top panels represent maximal projections of *fli1:nEGFP* stopped hearts, with blue dotted lines outlining the endocardium, and lower panels represent heat-maps of nucleus volume for the same embryonic heart. Unpaired two-tailed t-test. (N=5 embryos, 28hpf; N=5 embryos, 36hpf; N=5 embryos, 48hpf).

To overcome the difficulties associated with 3D manual segmentation, we developed a robust and quantitative deep-learning-based routine to obtain a reliable assessment of the cell volume from nuclei segmentation. Indeed, it has been reported that nuclear volume directly scales with the cell volume in different contexts (Cantwell & Nurse, 2019; Greiner et al., 2015; Huber & Gerace, 2007). We thus focused on extracting the nuclei volume which is easier to segment than the cell borders when in 3D. Quantitative data was obtained by using a trained classifier by deep learning. To confirm the validity of the approach, we showed that nucleus volume correlates with cell volume in our system (Figure 2.D). To demonstrate this correlation more clearly, we analyzed individual cells for both their cell surface and their nucleus volume and found a strong correlation between those two parameters (Figure 2.E).

We next assessed the temporal variation of cell volume during the progression of tissue convergence. At 28hpf, cells in the AVC were not different in size compared to cells located inside the ventricle (426.7±32.8 µm^3^ at 28hpf in the AVC), whereas they started to be smaller from 36hpf until 48hpf (390.6±12.9 µm^3^ at 36hpf, and of 284.9±6.1 µm^3^ at 48hpf) (Figure 2F). We confirmed these results by quantifying the mean distance between each nucleus and its three nearest neighbors (Figure S2.A). Moreover, we found that cell volume change is correlated with enrichment in actin filaments (F-actin) and phospho-myosin light chain (p-MLC) specifically in the cells that will form the AVC, suggesting that cells display active contractility in the AVC (Figure 1.D-E, Video S2). This observation indicates that the mechanical properties and behaviors of the cells in the AVC could be different from the endocardial cells located in the heart chambers. Overall, these results show that AVC EdCs undergo a substantial decrease in cell volume along with actin cortex remodeling. This argues that EdC contractility and cell volume decrease are involved in endocardial tissue remodelling.

### Cell volume decrease is independent of cell proliferation in the AVC

Differential cell proliferation rates or differential progression through the cell cycle between the three regions of the primitive heart tube (atrium, AVC, ventricle) could result in EdC cluster formation. We thus assessed if cell proliferation is associated with heart chamber-specific cell volume changes. To do so, we abolished cell division by using a combination of 30mM Hydroxyurea and 150µM Aphidicolin, two drugs that induce cell cycle arrest in the S phase (Figure 3.A). Embryos were treated from 30hpf until 48hpf without major phenotypic defects (Figure S1A-A’). Anti-phospho-Histone 3 immunolabeling confirmed that cell proliferation inhibition was effective in the presence of the drugs (Figure S3.B-B’). During the treatment, the number of EdCs doubled while cell numbers remained similar between 30hpf and 48hpf when proliferation was inhibited (Figure 3.A). The endocardial tissue and shape of the overall heart were qualitatively unchanged: the heart looped, and the overall volume of the heart was not significantly different between control and treated embryos (Figure 3.B, Figure S3.C-C’). When nuclei volume was quantified, we could not detect a difference in the cell size ratio between the three regions of the heart (atrium, AVC, ventricle) in controls and treated embryos (ratio of 1.48, 1.48, and 1.53 respectively in the atrium, AVC, and ventricle), suggesting a global scaling effect linking cell number and cell size (Figure 3.C). Similarly, we could not detect any difference in the mean nucleus-nucleus distance (ratio of 1.26, 1.27, and 1.15 respectively in the atrium, AVC, and ventricle) between controls and treated embryos (Figure S3.D). Importantly, cells were significantly smaller in the AVC of the treated embryos in the absence of proliferation comparable to cells located in the ventricle (Figure 3.C). These results show that cell volume decrease at the AVC is independent of EdC proliferation. Moreover, the nucleus-to-Golgi axis pattern at 48hpf was similar in treated and control embryos in both the ventricle (Figure 3.D) and the atrium (Figure 3.D’). This additionally suggests that tissue convergence occurs independently of cell proliferation.

**Figure 3.**
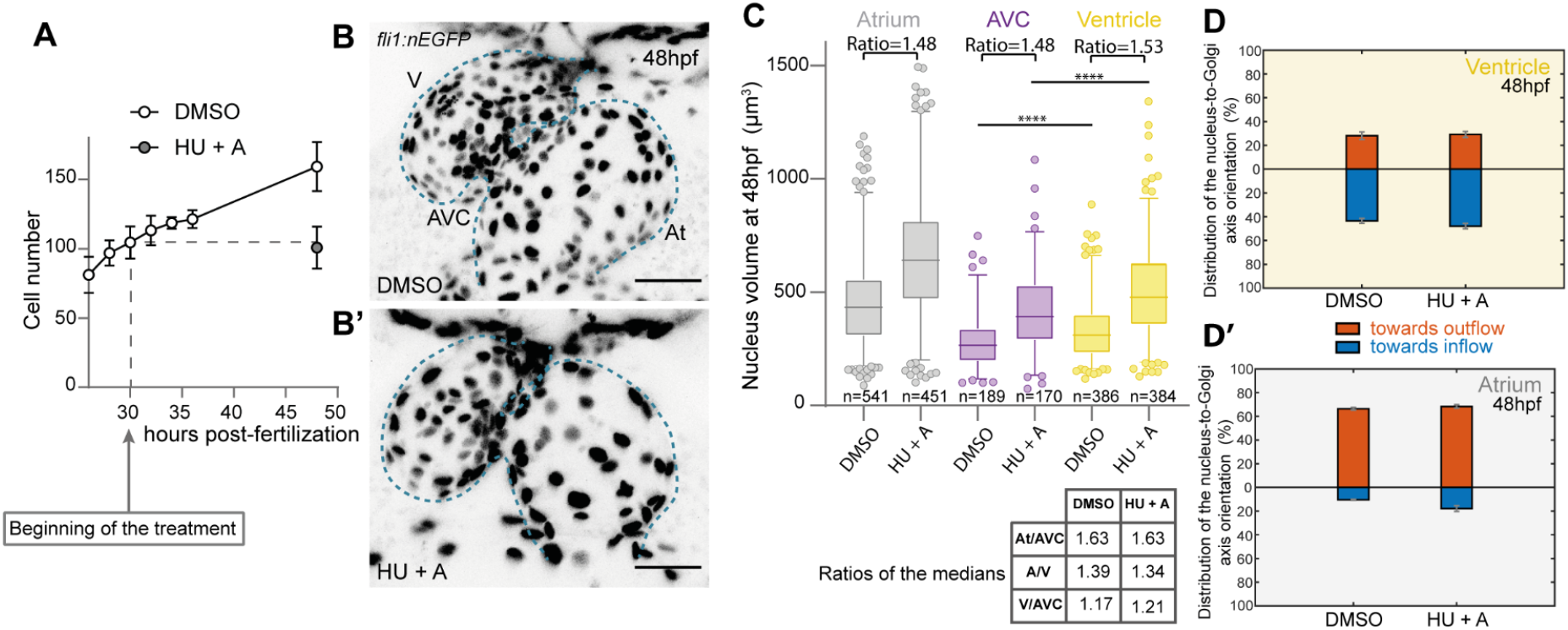
AVC cell volume reduction occurs independently of cell proliferation. (**A**) Total EdC number quantification in control embryos (treated with 0.5% DMSO), and embryos treated from 30hpf with a mix of two S-phase inhibitors, 30mM Hydroxyurea (HU) and 150 μM Aphidicolin (A). (for controls: N=5 embryos, 26hpf ; N=5 embryos, 28hpf ; N=5 embryos, 30hpf ; N=4 embryos, 34hpf ; N=5 embryos, 36hpf and N=7 embryos, 48hpf and N=10 HU+A embryos). Error bars represent the s.d (**B-B’**) Maximal projection of *fli1:nEGFP* embryos showing a control (DMSO) (**B**) and a treated embryo (HU+A) (**B’**) at 48 hpf. Blue dotted lines outline the endocardium. Scale bar, 50 μm. (**C**) Quantification of nucleus volume in atrium, AVC, and ventricle in controls (DMSO) and treated embryos (HU+A). The ratios of the medians are conserved between each region for both conditions. (N=7 embryos, DMSO and N=10 embryos, HU+A). Unpaired two-tailed t-test (**D-D’**) Quantification of the percentage of cells with the nucleus-to-Golgi axis towards the outflow and towards the inflow at 48hpf in controls (DMSO) and in treated embryos (HU+A) (N=3 embryos, DMSO and N=4 embryos, HU+A), for the ventricle (**D**) and for the atrium (**D’**). Error bars indicate the s.e.m.

### Mechanical forces are important regulators of cell volume decrease

As mechanical forces have been shown to be involved in endocardial tissue remodeling (Steed et al., 2016, Heckel et al., 2015, Dietrich et al., 2014), we next studied the impact of altering cardiac contraction and blood flow on EdC size. To do so, we first analyzed cell volume changes based on nuclei labelling in silent heart mutants (*sih*^*-/-*^), which carry a mutation in the gene *troponin T2a* (*tnnt2a)* and are therefore devoid of heart contraction and resulting blood flow (Sehnert et al., 2002) (Figure 4.A-A’). Cell volume did not show significant differences between the three regions of the heart (atrium, AVC, ventricle) and cells located in the AVC did not show a smaller volume compared to control embryos (nucleus volume of 481.7±25.5 µm^3^ in *sih*^-/-^, n=68 cells, N=12 embryos, nucleus volume of 273.2±7.6 µm^3^ in *sih*^+/+^ *sih*^*+/-*^, n=197 cells, N=8 embryos) (Figure 4.B). Less cells were present within the AVC compared to controls (Figure 4.D) and the nucleus-nucleus distance was significantly increased at the AVC in *sih*^*-/-*^ embryos (Figure S4.B). Interestingly, F-actin remodeling was absent in the AVC cells of the *sih*^*-/-*^ embryos compared to controls (Figure S4.A). Furthermore, at 48hpf cells located in the atrium showed random nucleus-to-Golgi axes distribution (Figure 4.F’) (towards the outflow: 35.3±2.9%; towards the inflow: 36.0±3.3% for n= 208 cells, N=6 embryos) different to what was observed in controls (towards the outflow: 68.7±1.4%, n=195 cells, N=3 embryos) (Figure 4.F’).Ventricular nucleus-to-Golgi axis reversal did not occur in *sih*^*-/-*^ embryos and instead cells tended to show a nucleus-to-Golgi axis towards the outflow (towards the outflow: 46.0±2.6%, towards the inflow:27.3±4.4%, n=314 cells, N=6 embryos) compared to controls (towards the inflow: 53.5±2.5%, n=217 cells, N=3 embryos) (Figure 4.F).These data are consistent with the observation that tissue convergence does not occur in absence of mechanical forces generated by the heartbeat (Boselli et al., 2017). To confirm these results, we partly inhibited heart activity by injecting a low concentration of the *tnnt2a* morpholino (Video S3) (Figure A’’). As expected, the flow velocity was lower (176.7±30.0 µm/s, N=12 embryos), compared to controls (1222.6±128.0 µm/s, N= 3 embryos) (Figure 4.E). Interestingly, cell volume decrease at the AVC was affected (nucleus volume of 360.9±11.7 µm^3^, n=153 cells, N= 12 embryos in *tnnt2a* morphants, nucleus volume of 324.9±10.5 µm^3^, n=114 cells, N=3 embryos) (Figure 4.C) suggesting that even subtle hemodynamic decrease alters cell volume. Overall, these results indicate that mechanical forces are important regulators of cell volume decrease.

**Figure 4.**
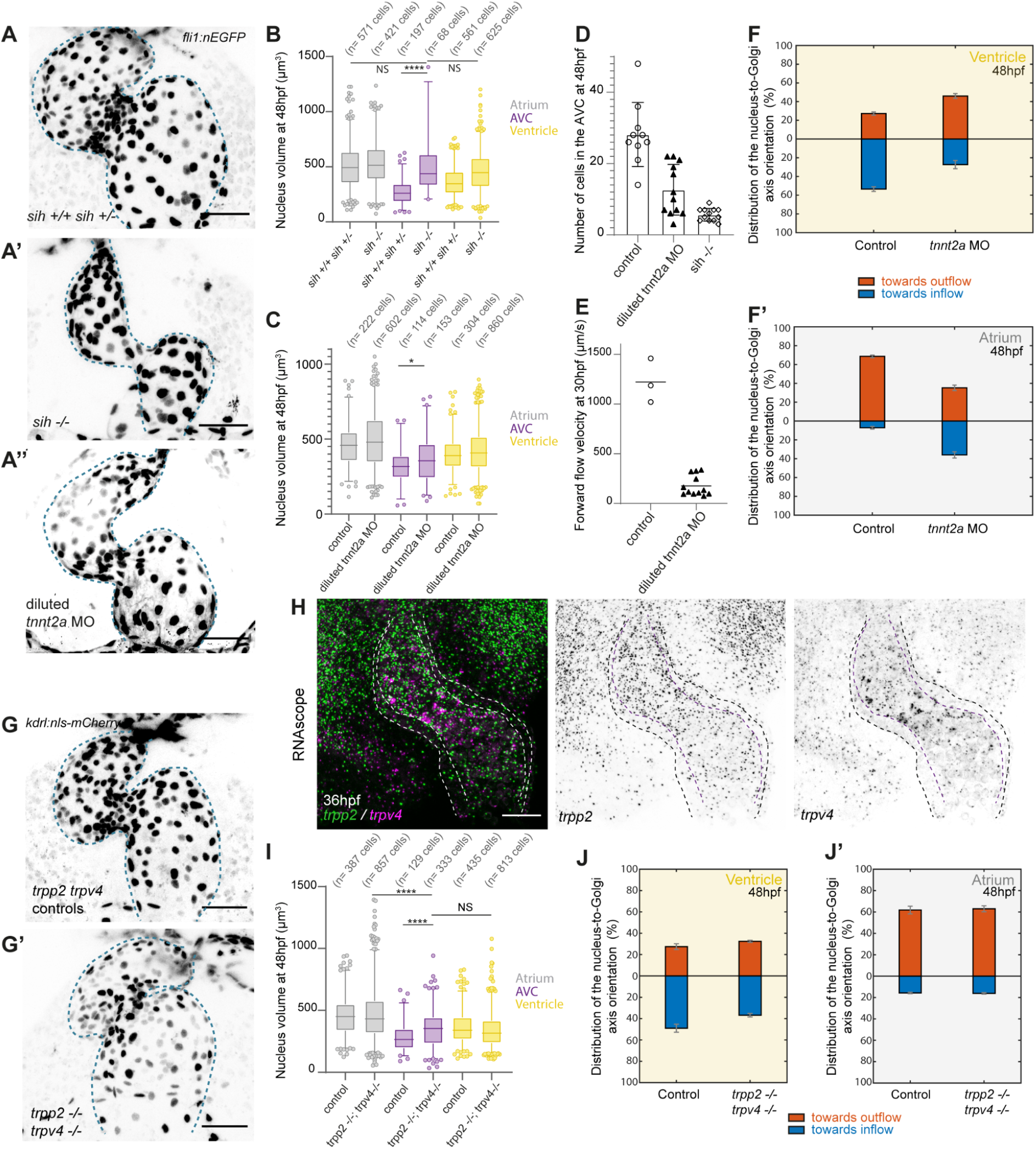
Mechanical forces and TRP channels mediate AVC endocardial cell volume decrease during early AVC morphogenesis. (**A-A’’**) Maximal projection of *kdrl :nls-mCherry* heart at 48 hpf for *sih*^*+/+*^ *sih*^*+/-*^ (**A**), *sih*^*-/-*^ (**A’**) and diluted *tnnt2a* morpholino (**A’**’). Scale bars, 50 μm. Blue dotted lines outline the endocardium (**B**) Quantification of the nucleus volume in atrium, AVC, and ventricle in both *sih*^*+/+*^ *sih*^*+/-*^ and *sih*^*-/-*^. (N= 8 *sih*^*+/+*^ *sih*^*+/-*^ embryos, N=12 *sih*^*-/-*^ embryos). Unpaired two-tailed t-test. (**C**) Quantification of the nucleus volume in the atrium, AVC, and ventricle in controls (sham) and diluted *tnnt2a* morpholino (MO)*-*injected embryos. (N= 3 controls (sham), N=12 embryos injected with diluted *tnnt2a* morpholino (MO)*-*injected embryos). Unpaired two-tailed t-test. (**D**) Number of EdC cells present in the AVC region based on the data presented in (A) for controls, diluted *tnnt2a* MO-injected embryos, and *sih*^*-/-*^. (**E**) Quantification of the forward flow (in the direction of the outflow) velocity at 30hpf based on manual tracking of fluorescent nano-droplets injected in the blood circulation, as explained in Figure (5.B). (**F-F’**) Quantification of the percentage of cells with the nucleus-to-Golgi axis towards the outflow and towards the inflow in the ventricle (**F**) and the atrium (**F’**) at 48hpf in controls (sham) and *tnnt2a* morphants (at high concentration). (N=3 embryos, sham and N=6 embryos, *tnnt2a* MO) Error bars indicate the s.e.m.(**G-G’**) Maximal projection of the stopped heart of *kdrl:nls-mCherry* embryos at 48hpf for *trpp2,trpv4* controls (**G**) (including trpp2^+/+^,trpv4^+/+^; *trpp2*^*+/-*^,*trpv4*^*+/+*^ and *trpp2*^*+/+*^,*trpv4*^*+/-*^) and their siblings *trpp2*^*-/-*^, *trpv4*^*-/-*^ (**G’**). Scale bars, 50 μm. Blue dotted lines outline the endocardium. (**H**) Maximum projection of confocal micrograph for *trpp2* and *trpv4* probes at 36hpf. The outer black dotted line represents the myocardial layer and the inner violet dotted line represents the endocardial layer (**I**) Quantification of nucleus volume in the atrium, AVC, and ventricle in *trpp2,trpv4* control embryos (including trpp2^+/+^,trpv4^+/+^; *trpp2*^*+/-*^,*trpv4*^*+/+*^ and *trpp2*^*+/+*^,*trpv4*^*+/-*^) and their *trpp2*^*-/-*^,*trpv4*^*-/-*^ siblings. (N= 15 embryos for *trpp2*^*-/-*^*trpv4*^*-/-*^ and N=6 control embryos). Unpaired two-tailed t-test. (**J-J’**) Quantification of the percentage of cells with the nucleus-to-Golgi axis towards the outflow and towards the inflow at 48hpf in controls and *trpp2-/- trpv4 -/-* in the ventricle (**J**) and in the atrium (**J’**). The antibody GM130 was used to label the golgi. (N= 3 embryos, controls and N= 4 embryos, *trpp2-/- trpv4 -/-*). Error bars represent the s.e.m.

### The stretch sensitive channels TRPP2 and TRPV4 are key modulators of cell volume decrease

The stretch-sensitive channels Transient Receptor Potential Polycystin 2 (TRPP2) and Transient Receptor Potential Vanilloid 4 (TRPV4) are important regulators of heart valve development (Heckel et al., 2015). Interestingly, TRPV4 is also a well-known contributor to cell volume regulation in astrocytes where it acts as an osmosensitive channel (Benfenati et al., 2011). We thus hypothesized that TRP channels could modulate cell volume during AVC morphogenesis. We investigated mutants for *trpv4* and *trpp2* channels. *trpp2* mRNA is ubiquitously distributed in the embryo but is enriched within the endocardial layer of the heart whereas *trpv4* is mostly expressed in the endocardium and specifically enriched in the AVC region (Figure 4.H). By looking at both *trpv4* ^*-/-*^ and *trpp2* ^*-/-*^ single mutant embryos, we found that EdC nucleus volume at the AVC was unchanged compared to controls (Figure S4.D). Since genetic compensation is a widespread feature in zebrafish (El-Brolosy & Stainier, 2017) we analysed double *trpv4* ^*-/-*^*;trpp2* ^-/-^ mutants (Figure 4.G-G’). Cell volume decrease was not observed in the AVC of *trpv4* ^*-/-*^*;trpp2* ^-/-^ compared to controls (nucleus volume of 351.9±7.9 µm^3^, n=333 cells, N=15 embryos in *trpv4* ^*-/-*^*;trpp2* ^-/-^, nucleus volume of 282.4±9.8 µm^3^, n=129 cells, N=6 in *trpv4* ^*+/+*^*;trpp2* ^+/+^) (Figure 4.I), and nucleus-nucleus distance was significantly increased in AVC EdCs of the double mutants (Figure S4.E). Importantly, heart rate was not significantly different between mutants, suggesting that heart function and flow forces are normal (Figure S4.F). These data show that the Trpp2 and Trpv4 mechanosensitive channels are important regulators of cell volume decrease. Moreover, the percentage of ventricular EdCs with nucleus-to-Golgi axis towards the inflow 36.9±1.7 % (n= 309 cells, N=4 embryos) is reduced compared to controls (49.0±3.8 % (n= 223 cells, N=3 embryos) (Figure 4.J) and the distribution of the nucleus-to-Golgi axes was unchanged in the atrium (Figure 4.J’). This suggests that the tissue convergence is affected in the absence of the TRP channels.

In the context of osmotically-driven cell volume increase (i.e. cell swelling) TRPV4 interacts with aquaporin channels to modulate cell volume (Benfenati et al., 2011; Conner et al., 2012; Iuso & Križaj, 2016). Aquaporins are passive transmembrane channels that can enhance membrane permeability (Ibata et al., 2011; Mola et al., 2016). We studied the spatio-temporal expression patterns of aquaporins and focused on two aquaporin channels present in the developing cardiovascular system: *aqp8a*.*1* and *aqp1a*.*1 (*Figure S2.B-S2.D). Both *aqp8a*.*1* and *aqp1a*.*1* mRNA are expressed within the heart (Figure S2.B), with expression starting from 30hpf specifically in AVC cells (9/32 embryos (*aqp8a*.*1*), 4/20 embryos (*aqp1a*.*1*)). At this stage, *aqp1a*.*1* is also expressed in the red blood cells, as previously reported (16/20 embryos) (Chen et al., 2010) (Figure S2.B). At 36hpf, *aqp8a*.*1* and *aqp1a*.*1* mRNAs are expressed in both AVC and OFT cells (24/27 embryos (*aqp8a*.*1*), 19/19 embryos (*aqp1a*.*1*). At 48 hpf, *aqp8a*.*1* is undetectable (36/36) whereas *aqp1a*.*1* mRNA expression is highly specific in the AVC and OFT regions (31/31). Using fluorescent probes, we confirm that these channels are expressed specifically in the AVC and OFT regions and further show that their expression is restricted to the endocardium (Figure S2.C). Interestingly, both the expression of *aqp8a*.*1* and *aqp1a*.*1* were absent in the *sih* ^*-/-*^ embryos, suggesting that their expression is dependent on mechanical forces (Figure S4.C). These data indicate that cells specifically located in the AVC are equipped with water channels whose expression depends on heart function.

### Hyaluronic acid modulates AVC endocardial cell volume changes

One of the most abundant components of the cardiac jelly is the glycosaminoglycan (GAG) hyaluronic acid (HA). GAG accumulation is known to apply osmotic pressure and to attract water (Cowman et al., 2015; Lockhart et al., 2011). *In vitro*, GAG leads to cell shrinkage of HEK cells (Joerges et al., 2012). HA is assembled by *hyaluronan synthase* genes and is then secreted into the ECM. In particular, *has2* is specifically expressed by EdCs of the AVC (Patra et al., 2011; Tong et al., 2014). As expected, *has2* expression in the heart starts at 30hpf and is restricted to the AVC from 30hpf to 48hpf (Figure S5.A). We used HA-binding protein (HA-BP) to study the protein localization (Figure 5.A). HA was found to be present exclusively within the cardiac jelly (Figure 5.A, Figure S5.B) and appears uniformly distributed throughout the cardiac jelly of the heart. We next assessed its potential role in cell volume regulation at the AVC by injecting hyaluronidase (HAase), which breaks down HA chains (Figure 5.B). HAase-injected embryos presented pericardial edema after 18 hours of treatment (Figure S5.C) but did not have heart rate defects at 48hpf (Figure S5.E). In this condition, HA was not detected inside the cardiac jelly of HAase-injected embryos, confirming that HA degradation was effective (Figure 5.C). Interestingly, F-actin remodelling in EdCs and cell volume decrease did not occur in the treated embryos (Figure S5.D) (N=7/7 HAase-injected embryos, N=7/7 controls). Cell volume decrease was not observed in the AVC of HAase-injected embryos compared to controls (nucleus volume of 422±10.0 µm^3^, n=177 cells, N=12 embryos in HAase-injected embryos, nucleus volume of 290.6±7.0 µm^3^, n= 266 cells, N=8 in controls) (Figure 5.E). Similarly, the AVC nuclei-nuclei distance was increased in HAase-injected embryos (Figure S5.F). Furthermore, the number of ventricular cells with nucleus-to-Golgi axis towards the inflow 30.1±2.2 % (n=523 cells, N=7 embryos) is reduced compared to controls 53.4±3.3% (n=425 cells, N=5 embryos) (Figure 5.F) and the distribution of the nucleus-to-Golgi axes was unchanged in the atrium (Figure 5.F’). This result indicates that tissue convergence is reduced in the absence of HA inside the cardiac jelly. We conclude that both mechanosensitivity and HA are essential modulators of the cell volume decrease in the AVC (Figure 5.G).

**Figure 5.**
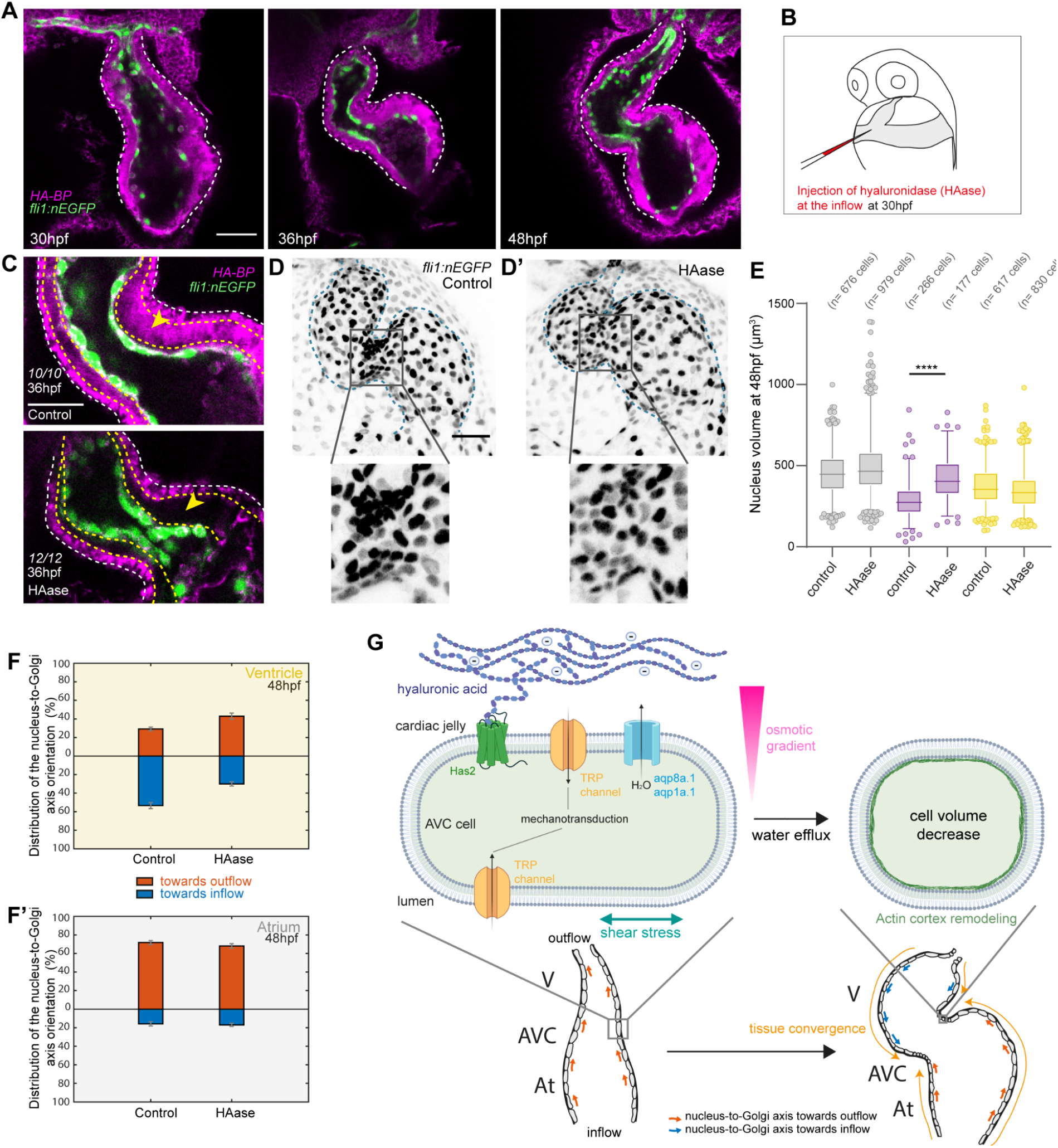
Hyaluronic acid located in the cardiac jelly modulates AVC endocardial cell volume changes. (**A**) Pictures of one z-plane showing the localization of HA-BP (magenta) by immunofluorescence on fixed and deyolked (to visualize the whole heart) *fli1:nEGFP* embryos (nuclei in green) at 30hpf (N=10 embryos), 36hpf (N=8 embryos), and 48hpf (N=12 embryos). Scale bar, 50 μm. White dotted lines outline the heart. (**B**) Schematic explaining the injection process of the hyaluronidase (HAase) inside the blood circulation, near the inflow region of the heart at 30hpf. (**C**) Pictures of one z-plane in the AVC region showing the presence of HA-BP inside the cardiac jelly (yellow arrow) for control (injected with PBS-injected) embryos (N=10/10) and the absence of HA-BP inside the cardiac jelly (yellow arrow) for HAase-injected embryos (N=12/12) at 36hpf. Scale bar, 30 μm. White dotted lines outline the heart. The yellow dotted lines mark the contours of the cardiac jelly. (**D-D’**) On the top panel: Maximal projections of stopped hearts at 48hpf of *fli:nEGFP* embryos for control (PBS-injected) embryos (**D**) and HAase-injected embryos (**D’**). On the bottom panel: zoom in the AVC region. Scale bar, 50 μm. Blue dotted lines outline the endocardium. (**E**) Quantification of the nucleus volume in the atrium, AVC, and ventricle in both controls (PBS-injected) embryos and HAase-injected embryos based on the raw data presented in (D). (N=8 controls, N=12 HAase-injected embryos). Two-tailed unpaired t-test. (**F-F’**) Quantification of the percentage of cells polarized towards the outflow and cells polarized towards the inflow in the ventricle (**F**) and the atrium (**F’**) at 48hpf in controls (PBS-injected) and HAase-injected embryos (N=5 embryos, controls, and N=7 embryos, HAase-injected) Error bars indicate the s.e.m. (**G**) Schematic illustrating the proposed mechanism for cell volume decrease at the AVC. Created partially with BioRender.com.

## DISCUSSION

Using *in vivo* imaging, we show that endocardial cell volume decrease is key for tissue convergence and AVC formation, setting the stage for subsequent valve formation (Pestel et al., 2016; Steed et al., 2016). We observed that cell volume decrease is concomitant with ventricular nucleus-to-Golgi axis reversal as well as F-actin remodeling in the AVC. Additionally, we show that cell volume change is independent of cell proliferation and is regulated by mechanical forces caused by heart function, TRP channels and HA located in the cardiac jelly. Together, our results show that cell volume decrease is an important cellular feature involved in cardiovascular morphogenesis in response to mechanotransduction activated by TRP channels. We propose that cell volume change may be a general cellular feature activated by mechanical forces to control tissus shape.

TRP channels are key players in the mechanotransduction pathway leading to heart valve formation (Duchemin et al., 2019; Heckel et al., 2015). The current model suggests that those channels sense oscillatory blood flow at the AVC, which is subsequently transduced into electrochemical information, leading to the expression of mechanosensitive genes (Steed, Boselli, et al., 2016, Heckel et al., 2015). Our results suggest that additional cues, on top of oscillatory blood flow, might affect the activity of the TRP channels through the generation of an osmotic pressure generated in the cardiac jelly. The components of the cardiac jelly are important for proper AVC formation (Derrick & Noël, 2021; Grassini et al., 2018; Hernandez et al., 2019). For example, modulating the expression of the *ugdh* gene encoding for the GAG building blocks or the alteration of HA deposition in the cardiac jelly leads to defects in AVC development (Segert et al., 2018; Walsh & Stainier, 2001). Here, we show that HA regulates the size of the EdCs in the AVC, revealed by the absence of a cell volume decrease in the hyaluronidase-treated embryos. Interestingly, it is known that GAG can establish an osmotic pressure (Cowman et al., 2015) and TRPV4 channels are potent osmosensitive channels (Hoffmann et al., 2009) that physically interact with aquaporin to modulate cell volume in response to osmotic stresses (Benfenati et al., 2011). Interestingly, the cells that lose volume were also found to be equipped with aquaporin channels (aqp8a.1, aqp1a.1). Considering that Aquaporin enhances water permeability (Ibata et al., 2011; Mola et al., 2016), the TRPV4-Aquaporin complex might increase the rate of water efflux following the establishment of an osmotic gradient. Moreover, the intracellular entry of calcium ions following the opening of the TRP channels might lead to the translocation of the aquaporin channels to the plasma membrane (Conner et al., 2012). With this, we propose a model where HA accumulation creates an osmotic pressure between the EdCs and the cardiac jelly to promote the cell volume decrease observed in the AVC (Figure 5.G). In this model, TRP channels would be involved both in the osmosensing as well as the shear stress sensing.

What is the driving force behind tissue convergence? At the tissue level, we observed a change of the nucleus-to-Golgi axis that is concomitant to cell volume changes and depending both on blood flow, and the presence of TRP channels and HA. As a consequence, the local cell volume decrease of EdCs in the AVC could pull on the surrounding cells and drive tissue convergence as well as the cell clustering. In addition, we cannot rule out that tissue convergence is in part due to an active migration of EdCs towards the AVC. Indeed, the presence of HA inside the ECM has been reported to be essential in the context of cell migration in several studies (Derrick & Noël, 2021) and ECM is key to regulate the turnover of angiogenic signals (De Angelis et al., 2017). Two main scenarios can therefore explain our results: active migration towards the AVC is taking place and then followed by an adaptive cell volume shrinkage and/or the cell volume shrinkage at the AVC pulls on the surrounding cells and leads to tissue convergence. Considering the small amplitude of the tissue movements, we favour the second scenario where cell volume decrease is sufficient to drive tissue movement as described in other species (Saias et al., 2015).

In summary, we observed that the loss of the mechanosensitive/osmosensitive TRP channels in the presence of normal blood flow affects the ability of AVC EdCs to decrease their cell volume and subsequent heart valve morphogenesis. These results indicate that EdCs process local mechanical signals to regulate their size through ion channels. The coordinated interactions between extracellular and intracellular processes involved in cell volume regulation are key to explain the mechanisms leading to tissue remodeling. Overall, a better understanding of these mechanisms will have implications for treating numerous pathological conditions including cardiovascular diseases such as congenital valvulopathies.

## METHODS

### Zebrafish husbandry, transgenic lines, morpholinos

Animal experiments were approved by the Animal Experimentation Committee of the Institutional Review Board of the IGBMC and followed ethical and animal welfare guidelines. 0.003% 1-phenyl-2-thiourea (PTU) (Sigma Aldrich, P7629) was added to 0.3X Danieau’s buffer at 8 hours post-fertilization in order to prevent pigment formation. Embryos were raised at 28.5 °C. The different zebrafish lines used in this study were: AB as wild-type line, *Tg(fli1a:B4GALT1galT-mCherry)*^*bns9*^ (Kwon et al., 2016), Tg*(fli1:nEGFP)*^*y7*^ (Roman et al., 2002), *Tg*(*ve-cad:ve-cadTS*) (Lagendijk et al., 2017), *Tg(fli1a:DsRed)* (Vatine et al., 2013); *Tg(fli1a:lifeact-EGFP)* (Phng et al., 2013), *Tg(kdrl:nls-mCherry)* (Nicenboim et al., 2015), *silent heart (sih)*^*tc300b*^ (Sehnert et al., 2002), *cup*^*tc321*^ (Schottenfeld et al., 2007), *trpv4*^*sa1671*^ (ZIRC). The genotyping primers were for *trpv4* the forward (5’-GCCTTTCAGCATGTTGTCCA-3’) and the reverse (5’-GGTTCCTGCTGGTCTACGTG-3’) primers with a Tm of 64.2°C for annealing and for *trpp2* the forward (5’-CCATTAGCCTGCACATTCAATC-3’) and the reverse (5’-ATCGCACTGCTCATCTGAAG-3’) primers with a Tm of 62.9°C for annealing. The *trpp2* homozygous mutant was selected phenotypically based on their curved tail phenotype. The morpholino (MO) used in this study targets *tnnt2a* (5’-CATGTTTGCTCTGATCTGACACGCA-3’) (GeneTools) (Sehnert et al., 2002). 5.8 ng of the *tnnt2a* MO was injected into the yolk at the one-cell stage for complete heartbeat arrest and 0.14 ng was injected to reduce heartbeat amplitude (referred to as diluted *tnnt2a* MO in text).

### Confocal imaging

For live imaging, dechorionated zebrafish embryos were anesthetized using 0.2mg.mL^-1^ of tricaine (Sigma Aldrich, A5040; stock at 8mg/mL adjusted to pH7-7.5) (to stop fish motion) or 50mM drug 2,3-butanedione monoxime (BDM) (Sigma Aldrich, B0753) for 10 minutes (to stop the heart) and then mounted in 1.2% UltraPure low melting-point agarose (Sigma Aldrich, 16520) that has the same concentration of tricaine or BDM as the media.

Confocal imaging was performed either on an up-right Leica SP8-Multiphoton confocal microscope (to image the stopped heart or fixed embryos) or on an inverted Leica spinning disk (to image the beating heart). The mounting was different on the two microscopes - a custom-designed mold was used for imaging on the up-right Leica SP8-MP confocal microscope (Chow et al., 2018) and a glass-bottom Petri dish (MateTek, P35G-0-14-C) for imaging on the inverted spinning disk microscope.

### Image analysis

#### Classification of cell polarity

Individual cell polarity was analysed by manually labelling individual nuclei and Golgi apparatus using the “spots” tool in the Imaris software (3D viewer). We then classified by eye the cell polarity of each cell based on the position of its Golgi apparatus relative to its cell nucleus: polarized towards the outflow, polarized towards the inflow, or no clear polarization.

#### F-actin intensity measurements

10 cells were analyzed for each region of the heart (atrium, AVC, ventricle) in *fli:LifeAct-eGFP* embryos. For each cell, 10 lines were drawn perpendicular to the plasma membrane with ImageJ and the maximal intensity was measured.

#### Cell volume estimation measurements

Measurement points were put with the Imaris software all around the cell membrane based on the VE-cadherin signal, *Tg*(*ve-cad:ve-cadTS*). The coordinates of these measurement points were inputted into Matlab, and the individual cell surface area was calculated using Delaunay triangulation-based surface reconstruction. Using the *Tg(fli1a:DsRed)* line to visualize the cytoplasm of endothelial cells, cell height measurements were made in Imaris software (3D viewer). Since AVC endocardial cells are cuboidal, an estimation of their cell volume was computed simply by multiplying the height of the cell with the cell surface area (Figure 2.B). For EdCs in the ventricle and atrium, the height of the cell was much higher at the part of the cell containing the cell nucleus (HN) than at other parts of the cell (HO). Thus, their cell volume was estimated by multiplying HN with the nucleus area, multiplying HO with the area of the cell not containing the nucleus, and adding the two products together (Figure 2.B).

#### Nuclei segmentation

Nucleus segmentation of 3D images was obtained with a Deep Learning based segmentation pipeline called StarDist3D (Weigert et al., 2020). The training of StarDist3D needs ground truth instance segmentation annotations which are cumbersome to obtain, if done fully manually. Hence, this manual annotation effort was partially circumvented by adopting an iterative approach for ground-truth annotation. Firstly, only one 3D image was fully manually annotated using the open-source Labkit plugin in Fiji (Schindelin et al., 2012). A StarDist3D network was then trained using the single annotated 3D stack. This trained network was then used to predict nucleus segmentations in a separate 3D image. These predictions were then manually curated such that all segmentation errors were removed. The curated image, together with the first, manually annotated image, were then used to train yet another StarDist3D network, and this iterative training, prediction, curation loop can be continued.

In this work, this process was iterated 4 times, leading to a final StarDist3D network trained on a total of 5 full 3D images. While manual annotation of the first ground truth image took roughly 4 hours, the curation times for the subsequent 4 iterations reduced to ∼2 hours, ∼1 hour, ∼30 minutes and ∼10 minutes, respectively. Network training was performed using patches of size 48×96×96 and a batch size of 2 for 400 epochs and 100 steps per epoch, using the default parameters of the public implementation of StarDist3D (https://github.com/stardist/stardist).

The final network was used to obtain instance segmentations for all datasets used in this work. After segmentation, a custom python script was used to compute volume and 3D position of each segmented cell nucleus. By matching the 3D coordinates of nuclei positions back to the original image, nuclei were manually sorted into three groups (atrium, AVC, and ventricle) based on the region of the heart they are located in.

To obtain the mean distance between each cell and its three nearest neighbours, the 3D coordinates of the segmented nuclei were inputted into a custom Matlab script. For each nucleus, the Matlab script calculates the distance between the nucleus and all the other nuclei within the heart. Then, those values are sorted according to size, and the mean value for the three shortest distances was calculated.

### Immunofluorescence

Zebrafish embryos were dechorionated and fixed for 2-3 hours at room temperature (RT) at the desired developmental stage in either 4% paraformaldehyde (PFA) in 1X phosphate-buffered saline solution (PBS) or in 4g of PFA diluted in 100mL of Fish Fix Buffer (1L: 1X PBS, 120µL 1M CaCl2, 40g sucrose) for easy removal of the yolk with forceps. At early developmental stages, the removal of the yolk was beneficial for imaging the whole endocardial tissue, in particular the part of the endocardium near the outflow tract. Therefore, the yolk was removed for cell polarity, HA-BP immunofluorescence, and RNAscope experiments.

After fixation, embryos were washed in PBS implemented with 0.1% Tween20 (PBST), three times for 5 minutes. For GM130, anti-phosphorylated histone H3 antibody, and phalloidin stainings, embryos were permeabilized in 1X PBS containing 0.1% Triton X-100 (PBST) (Sigma-Aldrich, T8787) (GM130 antibody) or 0.5% Triton X-100 (phospho-H3 antibody), overnight at 4°C. Embryos were then incubated in blocking solution (PBS with 0.1% Triton X-100, 2% BSA (H2B, 1005-70), 5% NGS (Coger, VS-1000) (GM130) (Sepich & Solnica-Krezel, 2016) or (PBS with 0.5% Triton X-100, 2 mg/mL BSA, 2% NGS) overnight at 4°C. Primary antibodies were then added at the following dilutions: Mouse-anti-GM130, 1:100, BD Bioscicence-610822), (Rabbit-anti-PH3, 1:100, Millipore-06-570 in fresh blocking buffer for 2 days. Then, embryos were washed over 4 hours (with solution changes every 30 minutes) in PBST. Secondary antibodies were added: Goat-anti-Mouse (Alexa Fluor 647, Invitrogen, A32728) or Goat-anti-Rabbit (Alexa Fluor 594, Invitrogen, A11037), and counterstained with Phalloidin when necessary (Alexa Fluor 568, 1:50, Invitrogen, A12380).

For Hyaluronic Acid-Binding Protein (HA-BP, Sigma-Aldrich, 385911) immunofluorescence, the experimental procedure described in (Munjal et al., 2020) was performed on deyolked embryos.

### Pharmacological treatments

#### Hydroxyurea and Aphidicolin

Larvae were incubated from 30hpf to 48hpf with a mix of two S-phase inhibitors: 30 mM Hydroxyurea (HU) (Sigma-Aldrich, H8627; diluted in water) and 150 μM Aphidicolin (A) (Sigma-Aldrich, 89458; diluted in DMSO).

#### Hyaluronidase pharmacological treatment

Zebrafish embryos were dechorionated, anesthetized with 0.2mg.mL^-^1 of tricaine, and mounted on a glass-bottom Petri dish (MateTek, P35G-0-14-C) with 1.2% UltraPure low melting-point agarose (Sigma Aldrich, 16520). The injection solution was prepared as follows: 1X PBS with 0.5% Phenol Red (Sigma Aldrich, P0290) for controls and Hyaluronidase from *Streptomyces Hyalurolyticus* (Sigma-Aldrich, H1136, diluted in 1X PBS) with 0.5% Phenol Red for treated embryos. The solutions were injected into the embryos via the cardinal vein at 30 hpf near the inflow region of the heart using glass capillaries and a NanoInjectII injector (Drummond Scientific, Broomall, PA, USA). Embryos were then carefully removed from agarose using forceps before returning them to 0.3X Danieau, 0.003% PTU and placed in a 28.5°C incubator.

### Flow analysis

Embryos were injected as described in the section above, except here the injection mix was made by diluting a solution containing 95nm 561nm fluorescent nano-droplets with 3.3% Dil-TPB suspension (Kilin et al., 2014) 1:1000 in PBS. Embryos were removed carefully from the low melting point agarose and mounted again in order to be imaged under the spinning disk microscope with a Leica 40X (NA 1.1) water immersion objective. The analysis of the flow was then realized with the Manual Tracking plugin available under FiJi.

### In situ hybridization

Whole-mount in situ hybridization (ISH) was performed as in Thisse and Thisse (2008). Aqp8a.1 and aqp1a.1 probes were generated for this study. The *aqp8a*.*1* probe was generated with a fully sequence cDNA clone (IRBOp991A0150D – Source Bioscience) with the primers: forward primer containing the T7 promoter (5’-TAATACGACTCACTATAGCTGAAGCTCCGGGCAG-3’) and the reverse primer containing the Sp6 promoter (5’ ATTTAGGTGACACTATAGCCTCTTCAGTTCCTTCTTCCATC-3’).The aqp1a.1 probe was generated from whole extracted cDNA at 30hpf and amplified with the primers (5’-GTCATGAACGAGCTGAAGAGC-3’) and (5’-GGGTCACTTTGAGGACATCTC-3’),incorporated into the pCR-BluntII-TOPO vector (Invitrogen, 45-0245) (containing both the SP6 and T7 promoters), linearized with NotI. Both were then subsequently transcribed with the SP6 enzyme (mMessage mMachine SP6 transcription kit (Ambion)) in order to obtain the antisense RNA probe that was then purified with the RNeasy kit (Qiagen – 74104).

The has2 probe was generated from the plasmid PBSK-dg42II containing cDNA of the zebrafish has2 (provided by the Bakkers lab, The Netherlands), linearized with XbaI and subsequently transcribed using the T7 polymerase.

Imaging of ISH was then realized using a Leica M165 macroscope with a TrueChrome Metrics (Tucsen) with a Leica 1.0X objective (10450028).

### RNAscope

RNAscope experiments were performed using the RNAscope Fluorescent Multiplex kit (Advanced Cell Diagnostics, 323110) and by following the manufacturer’s guidelines.

### Statistical analysis

For the statistical analysis of the data, Student’s paired t-test with a two-tailed distribution or Mann-Whitney test with a two-tailed distribution were performed using the Prism software. For *p* values: < 0.05 ^*^; < 0.01 ^**^, < 0.001 ^***^, < 0.0001 ^****^

In each figure, the statistical test performed on the data, the number of analyzed cells (n) and/or the number of embryos (N), as well as the meaning of error bars is stated. Box plots were generated with the Prism software where horizontal lines show the median. The whiskers extend to the 2.5th and 97.5th percentiles. Data points outside whiskers are shown as individual circles.

## ACKNOWLEDGMENTS

We thank R. Chow as well as D. Riveline, S. Quintin and the participants of the “Biophysics club” at the IGBMC for the fruitful discussions and help with the manuscript. We also thank the members of the Norden lab for their feedback on the project. We are grateful to all the staff members of the imaging platform at IGBMC, especially E. Grandgirard and also the members of the IGBMC fish facility (S. Pajot, S. Geschier and C. Moebs). This project has received funding from the European Research Council (ERC) under the European Union’s Horizon 2020 research and innovation programme: GA N°682939, Agence Nationale de la Recherche: ANR-15-CE13-0015-01, ANR-10-IDEX-0002–02, ANR-12-ISV2-0001-01 and ANR-10-LABX-0030-INRT and the European Molecular Biology Organization Young Investigator Program. HV was supported by the IGBMC International PhD program: ANR-10-LABX-0030-INRT.

## AUTHOR CONTRIBUTIONS

Conceptualization, H.V. and J.V.; Visualization, H.V; Methodology, H.V., M.P., C.N., and F.J.; Resources, J.V.; Data Curation, H.V., C.V-P. M.P; Formal Analysis, H.V. and J.V.; Validation, H.V., C.V-P and J.V.; Writing– Original Draft, H.V and J.V.; Funding Acquisition J.V.; Supervision, J.V.; Project Administration, J.V.

## COMPETING INTERESTS

The authors declare no competing or financial interests.

**Figure S1.**
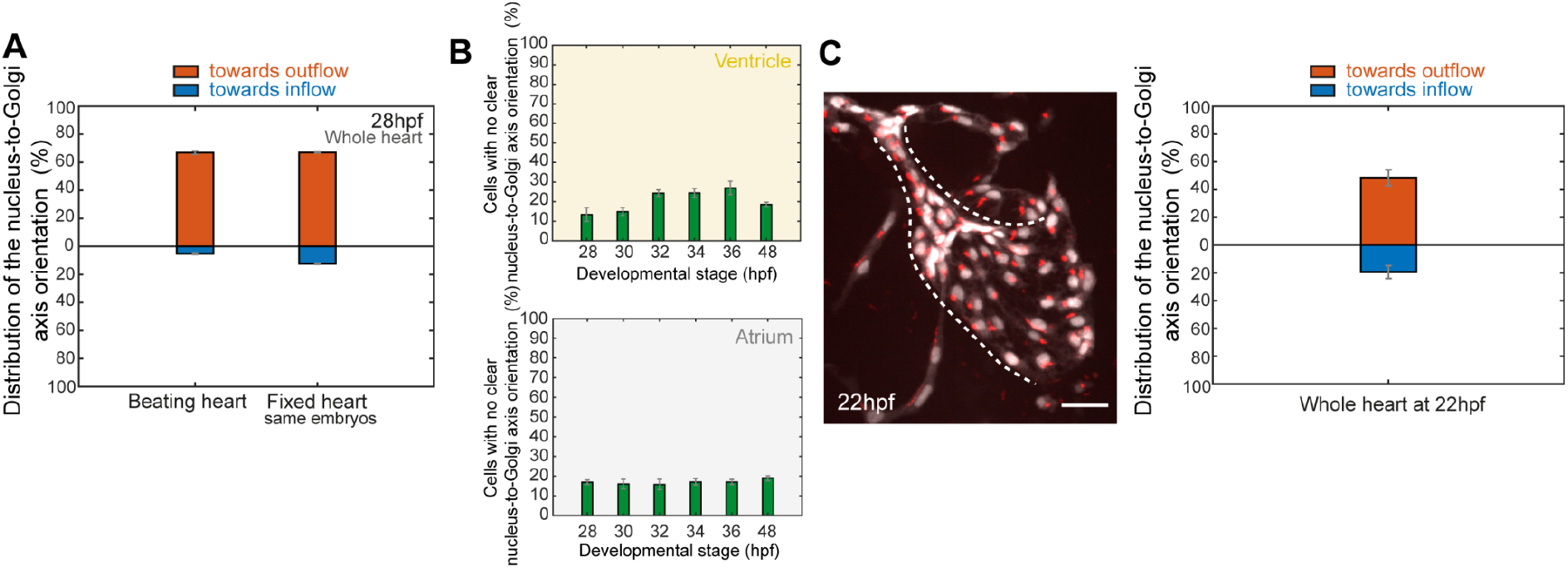
(**A**) Quantification of the percentage of EdCs nucleus-to-Golgi axis towards the outflow and towards the inflow at 28hpf in hearts that are beating and in the same hearts after fixation, showing that fixation does not affect the cell orientation pattern within the endocardium (N=3/3 embryos). (**B**) Quantification of the percentage of cells with no clear orientation of the nucleus-to-Golgi axis from 28hpf to 48hpf, in both ventricle and atrium. (N=5 embryos, 28hpf ; N=5 embryos, 30hpf ; N=4 embryos, 32hpf ; N=4 embryos, 34hpf ; N=5 embryos, 36hpf ; N=7 embryos, 48 hpf). Error bars show the s.e.m (**C**) Maximal projection of a fixed and deyolked heart at 22hpf (Scale bar, 30 μm.) and analysis of the EdC nucleus-to-Golgi axis distributions within the whole heart tube (N= 3 embryos).

**Figure S2.**
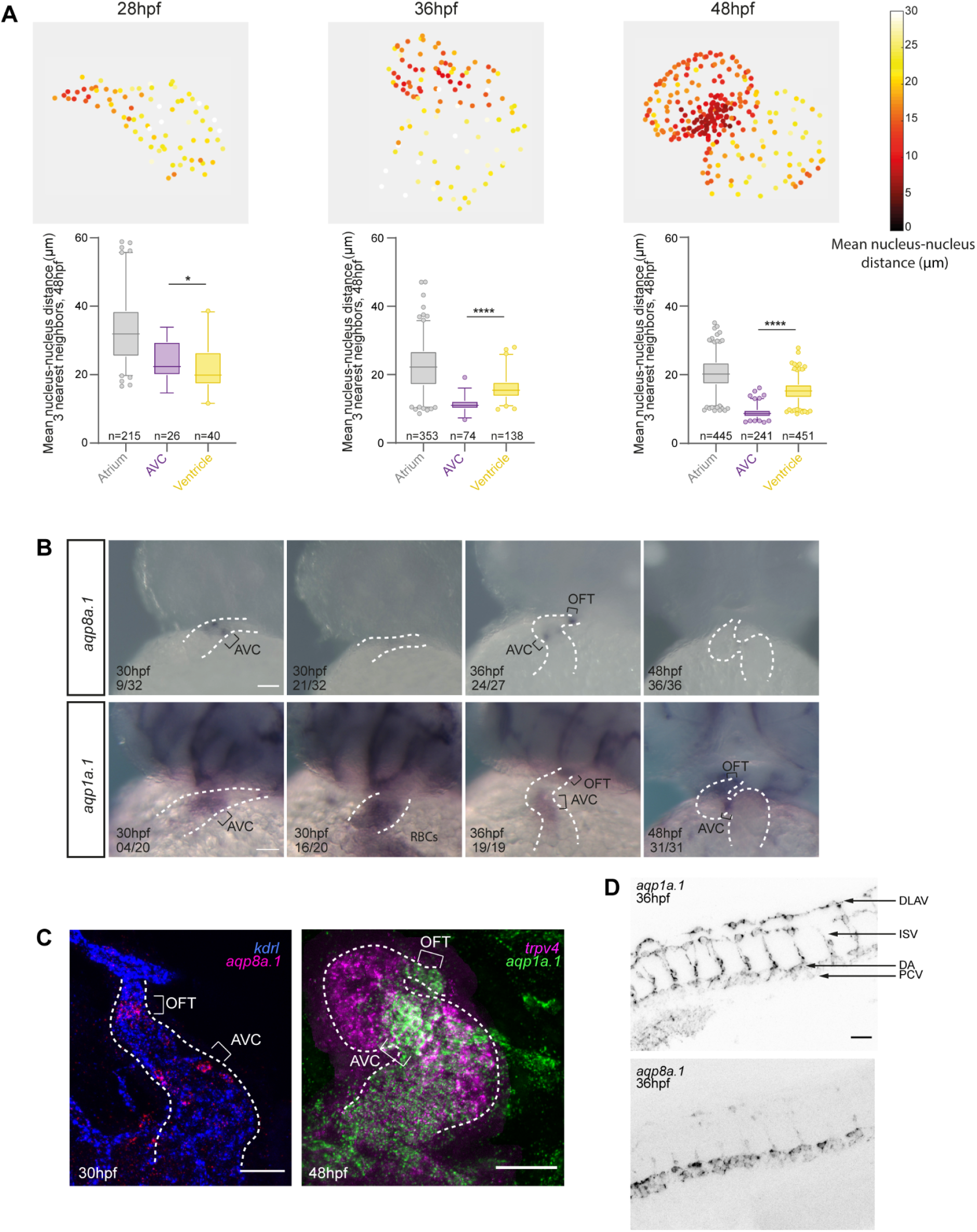
(**A**) Heat-maps of the distance between a nucleus and its three nearest neighbours (top panel) and quantifications of those distances by heart region (Atrium, AVC, ventricle) at 28hpf, 36hpf, and 48hpf. (N=5 embryos, 28hpf; N=5 embryos, 36hpf; N=5 embryos, 48hpf). (**B**) *In situ* hybridization with *aqp8a*.*1*and *aqp1a*.*1* probes in AB embryos at 30hpf, 36hpf, and 48hpf. White dotted lines outline the heart. Scale bars, 50 μm. (**C**) *aqp8a*.*1* and *aqp1a*.*1* mRNA expression profile in the endocardium. On the left: maximal projection of a multiplex RNAscope with the *kdrl* and *aqp8a*.*1* probes at 30hpf (N=7/7 embryos). On the right: maximal projection of a multiplex RNAscope using *trpv4* and *aqp1a*.*1* probes at 48hpf (N=5/5 embryos). Scale bars, 50 μm. (**D**) *aqp8a*.*1* and *aqp1a*.*1* mRNA expression in the endothelium of the cardiovascular system. DLAV= dorsal longitudinal anastomotic vessels, ISV= Intersegmental vessels, DA= dorsal aorta, PCV= posterior cardinal vein. Scale bars, 50 μm.

**Figure S3.**
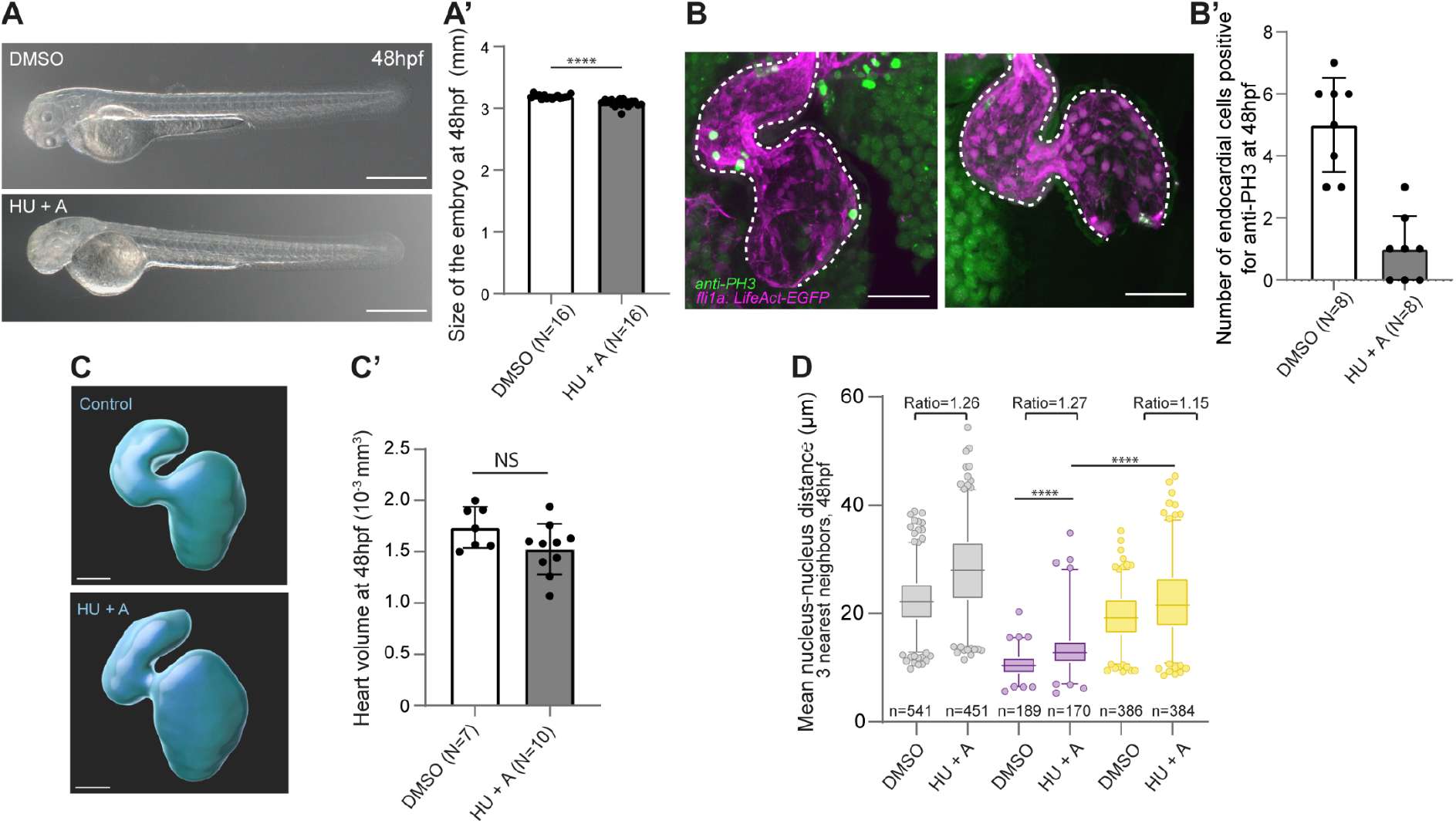
(**A**) Bright-field images of the embryo phenotype at 48hpf for controls (embryos treated with 0.5% DMSO from 30hpf to 48hpf) and treated embryos (embryos treated with 30mM Hydroxyurea (HU) and 150 μM Aphidicolin (A) from 30hpf to 48hpf). Scale bars, 0.5mm (**A’**) Measurements of total embryo size based on bright-field images at 48hpf (N=16 embryos, DMSO and N=16 embryos, HU+A). Mann-Whitney test. Error bars represent the s.d. (**B**) Immunofluorescence with anti-phosphorylated histone H3 in controls (DMSO) and treated (HU+A) in *fli1a:LifeAct-eGFP* embryos. Scale bars, 50 μm. White dotted lines outline the endocardium (**B’**) Analysis based on the immunofluorescence data in (B) showing the number of EdCs in the heart with anti-PH3 antibody signal at 48hpf (N=8 embryos, DMSO and N=8 embryos, HU+A). Bar plot with the mean and error bars representing the s.d. (**C**) Imaris 3D viewer representation of the whole heart volume at 48hpf. Scale bars, 50 μm. (**C’**) Quantification of the heart volume based on the Imaris data in (C) (N=7 embryos, DMSO and N=10 embryos, HU+A). Bar plot with the mean and the error bars represent the s.d. Two-tailed Mann-Whitney test. (**D**) Quantitative analysis of the mean distance between one nucleus and its three nearest neighbours at 48hpf for controls (DMSO) and treated (HU+A) *fli1:nEGFP* embryos.

**Figure S4.**
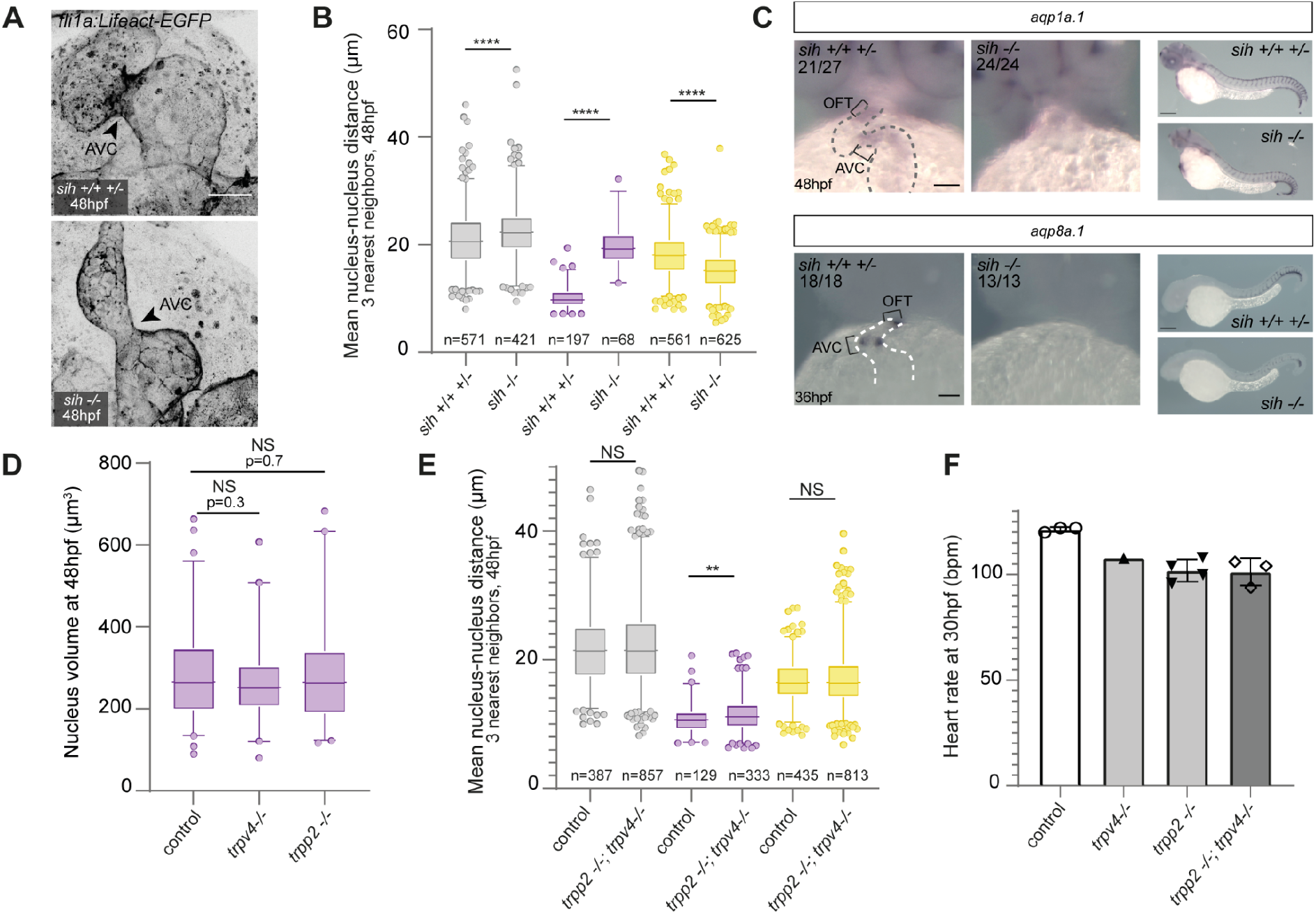
(**A**) Maximal projections of stopped hearts for *sih*^*+/+*^ *sih*^*+/-*^; *fli1a:LifeAct-EGFP* showing enrichment of F-actin signal at the AVC (arrowheads) and for their siblings *sih*^*-/-*^; *fli1a :LifeAct-EGFP* showing the absence of F-actin signal at the AVC (arrowheads). (**B**) Quantitative analysis of the mean distance between one nucleus and its three nearest neighbours at 48hpf for *sih*^*+/+*^ *sih*^*+/-*^ and *sih*^*-/-*^ *kdrl:nls-mCherry* embryos, based on the raw data presented in Figure 4.A (N= 8 *sih*^*+/+*^ *sih*^*+/-*^ embryos, N=12 *sih*^*-/-*^ embryos). Unpaired two-tailed t-test. (**C**) In situ hybridizations with the aqp8a.1 and aqp1a.1 probes in *sih*^*+/+*^, *sih*^*+/-*^ and *sih*^*-/-*^. Scale bars, 50 μm for the heart zoom and 200 μm for the entire embryo. Gray or white dotted lines outline the heart. (**D**) Quantification of the nucleus volume in the AVC in both *trpp2,trpv4* controls (including *trpp2*^*+/+*^,*trpv4*^*+/+*^; *trpp2*^*+/-*^,*trpv4*^*+/+*^ and *trpp2*^*+/+*^,*trpv4*^*+/-*^) and their *trpp2*^*-/-*^ (*trpp2*^*-/-*^,*trpv4*^*+/+*^) and *trpv4*^*-/-*^ (including *trpp2*^*+/+*^,*trpv4*^*-/-*^ and *trpp2*^*+/-*^,*trpv4*^*-/-*^) siblings. (N=6 embryos for the controls, N=4 embryos for *trpv4*^*-/-*^, N=5 embryos for *trpp2*^*-/-*^). Unpaired two-tailed t-test. (**E**) Quantitative analysis of the mean distance between one nucleus and its three nearest neighbours at 48hpf for *trpp2, trpv4* controls (including trpp2^+/+^,trpv4^+/+^; *trpp2*^*+/-*^,*trpv4*^*+/+*^ and *trpp2*^*+/+*^,*trpv4*^*+/-*^) and their siblings *trpp2*^*-/-*^,*trpv4*^*-/-*^ embryos. (N= 15 embryos for *trpp2*^*-/-*^ *trpv4*^*-/-*^ and N=6 control embryos). Error bars represent the s.e.m (**F**) Quantification of the heart rate (in bpm: beats per minute) at 30hpf at room temperature in controls (including trpp2^+/+^,trpv4^+/+^; *trpp2*^*+/-*^,*trpv4*^*+/+*^ and *trpp2*^*+/+*^,*trpv4*^*+/-*^), *trpp2* single mutant, *trpv4* single mutant and *trpv4 trpp2* double mutants. Bar plots represent the mean and the error bars represent the s.d.

**Figure S5.**
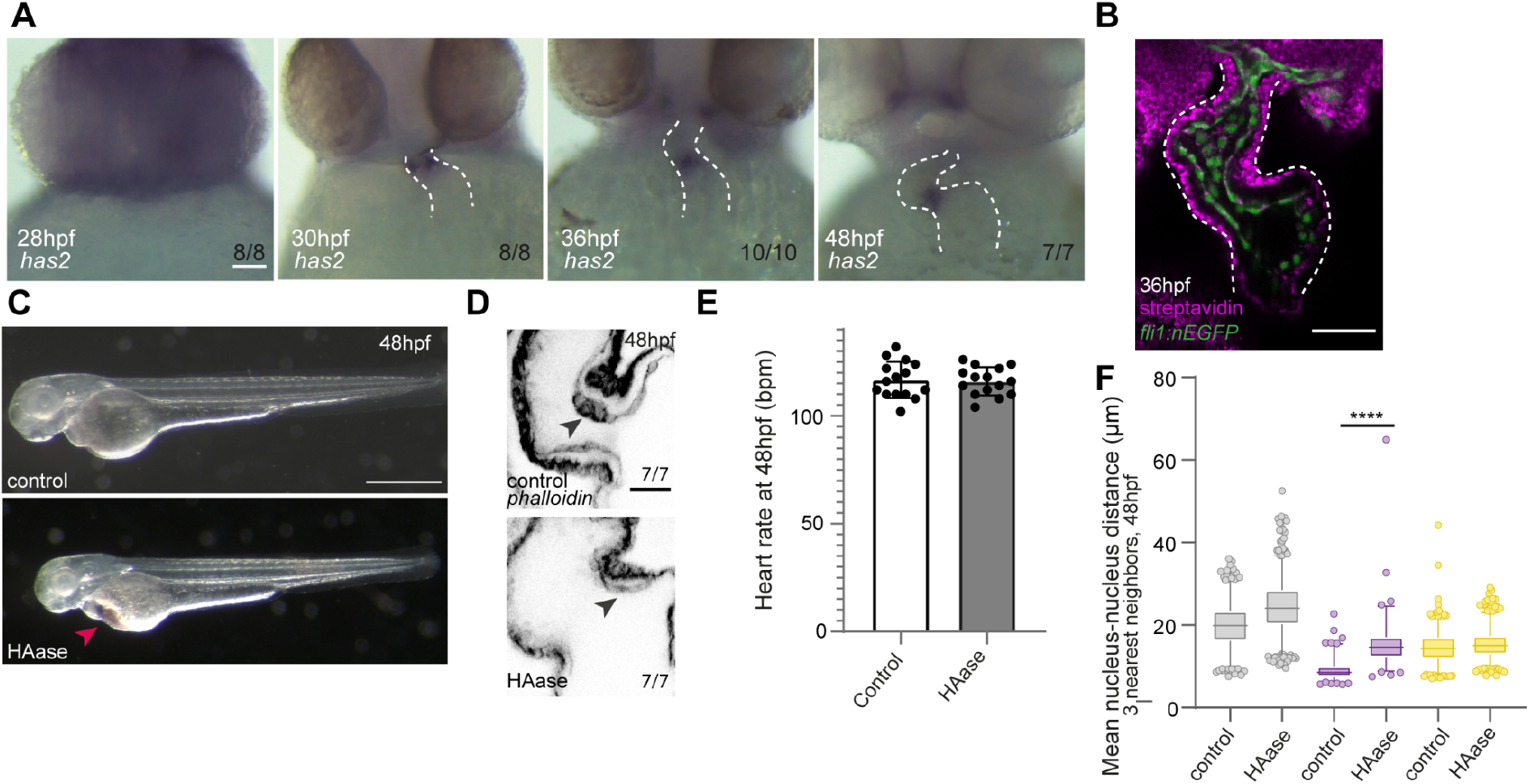
(**A**) *In situ* hybridization images with *has2* probe in AB embryos at 28hpf, 30hpf, 36hpf, and 48hpf. Scale bar, 50 μm. White dotted lines outline the heart. (**B**) Control immunofluorescence where Streptavidin was added in the absence of HA-BP resulting in a strong background signal in myocardial cells and a weaker background signal in endocardial cells. (N=6/6 embryos). Scale bar,50 μm. White dotted lines outline the endocardium. (**C**) Phenotype of the embryo at 48hpf. The embryos injected with HAase at 30hpf present edema (red arrow). Scale bar, 0.5mm (**D**) Phalloidin stainings in controls (N=7 embryos) and HAase-injected embryos (N=7 embryos). Black arrows point towards the endocardial cells of the AVC. (**E**) Quantification of the heart rate (in beats per minute) at 48hpf (N=15 controls, N=15 HAase-injected embryos) based on counting the number of beats per minute at room temperature. Bar plot with the mean and the error bars represent the s.d. (**F**) Quantitative analysis of the mean distance between one nucleus and its three nearest neighbours at 48hpf for controls (PBS-injected) embryos and HAase-injected embryos. Two-tailed unpaired t-test.

## Supplementary videos legends

**<u>Video S1:</u>**

Related to Figure 1 (A). Reconstructed 3D beating heart of a *fli1a: B4GALT1-mCherry; fli1: nEGFP* embryo (in which the nucleus and Golgi are simultaneously labelled) at 48hpf. The white arrows present on stopped frames of the video represent the nucleus-to-Golgi axis orientation which is mainly towards the outflow for the atrial chamber, and mainly towards the inflow for the ventricular chamber.

**<u>Video S2:</u>**

Related to Figure 1 (D). Reconstructed 3D beating heart of a *fli:LifeAct-eGFP* embryo at 48hpf. The video shows the accumulation of the F-actin signal within the AVC region.

**<u>Video S3:</u>**

Bright-field video of the way the heart is beating at 48hpf in a control embryo (on the left) and in an embryo injected with a low amount of the *tnnt2a* morpholino (on the right).

